## Supplemental Table 1 for "A mating-induced reproductive gene promotes *Anopheles* tolerance to *Plasmodium falciparum* infection"

**S1 Table**

| Figure | Response variable | Model | Effect Test Outputs |
|---|---|---|---|
| **1A** | egg number | GLMM, Gaussian distribution | mating: LRT $X^2_1$ = 0.009, $p$ = 0.93<br>infection: LRT $X^2_1$ = 0.76, $p$ = 0.38<br>mating*infection: LRT $X^2_1$ = 3.1, $p$ = 0.08 |
| **1B** | egg prevalence | GLMM, binomial distribution | **mating: LRT $X^2_1$ = 16.26, $p$ < 0.001**<br>infection: LRT $X^2_1$ = 3.24, $p$ = 0.07<br>mating*infection: LRT $X^2_1$ = 0.0001, $p$ = 0.99 |
| | egg number | GLMM, Gaussian distribution | **mating: LRT $X^2_1$ = 20.22, $p$ < 0.001**<br>infection: LRT $X^2_1$ = 0.007, $p$ = 0.94<br>mating*infection: LRT $X^2_1$ = 0.99, $p$ = 0.32 |
| **2A** | oocyst prevalence | GLMM, binomial distribution | mating: LRT $X^2_1$ = 1.08, $p$ = 0.30 |
| | oocyst intensity | GLMM, zero-truncated negative binomial distribution | mating: LRT $X^2_1$ = 0, $p$ = 1.00 |
| **2B** | oocyst prevalence | GLMM, binomial distribution | mating: LRT $X^2_1$ = 0.007, $p$ = 0.93 |
| | oocyst intensity | GLMM, zero-truncated negative binomial distribution | mating: LRT $X^2_1$ = 1.47, $p$ = 0.23 |
| **3A** | egg number | GLMM, zero-inflated negative binomial distribution | treatment: LRT $X^2_1$ = 0.16, $p$ = 0.69<br>gametocytemia: LRT $X^2_3$ = 6.33, $p$ = 0.10<br>**treatment*gametocytemia: LRT $X^2_3$ = 16.15, $p$ = 0.001** |
| **3B** | oocyst prevalence | GLMM, binomial distribution | treatment: LRT $X^2_1$ = 0.60, $p$ = 0.44<br>gametocytemia: LRT $X^2_3$ = 5.78, $p$ = 0.12<br>treatment*gametocytemia: LRT $X^2_3$ = 4.63, $p$ = 0.20 |
| | oocyst intensity | GLMM, zero-truncated negative binomial distribution | treatment: LRT $X^2_1$ = 0.85, $p$ = 0.36<br>**gametocytemia: LRT $X^2_3$ = 20.36, $p$ < 0.001**<br>treatment*gametocytemia: LRT $X^2_3$ = 0.85, $p$ = 0.84 |
| **3C** | egg number | GLMM, zero-inflated negative binomial distribution | treatment: LRT $X^2_1$ = 0.08, $p$ = 0.78<br>oocyst number: LRT $X^2_1$ = 1.89, $p$ = 0.17<br>**treatment*oocyst number: LRT $X^2_1$ = 7.77, $p$ = 0.005** |
| **3D** | egg prevalence | GLMM, binomial distribution | treatment: LRT $X^2_1$ = 0.06, $p$ = 0.81<br>oocyst number: LRT $X^2_1$ = 1.07, $p$ = 0.30<br>**treatment*oocyst number: LRT $X^2_1$ = 7.18, $p$ = 0.007** |
