## Supplemental Table 2 for "A mating-induced reproductive gene promotes *Anopheles* tolerance to *Plasmodium falciparum* infection"

**S2 Table**

| Donor | Gametocytes (/µl of blood) | dsRNA | Sample size | Oocyst intensity range | Oocyst intensity mean | Oocyst intensity median | Prevalence of infection (%) |
|---|---|---|---|---|---|---|---|
| 1 | 104 | ds*Control* | 13 | 0-15 | 6.1 | 5.0 | 69.2 |
| | | ds*MISO* | 31 | 0-22 | 5.5 | 3.0 | 71.0 |
| 2 | 304 | ds*Control* | 34 | 0-178 | 30.9 | 16.0 | 76.5 |
| | | ds*MISO* | 24 | 0-127 | 33.5 | 17.5 | 75.0 |
| 3 | 72 | ds*Control* | 19 | 0-22 | 4.2 | 2.0 | 73.7 |
| | | ds*MISO* | 20 | 0-30 | 8.5 | 5.0 | 95.0 |
| 4 | 128 | ds*Control* | 22 | 0-87 | 26.3 | 27.0 | 90.9 |
| | | ds*MISO* | 18 | 0-62 | 27.2 | 24.5 | 83.3 |
| 5 | 72 | ds*Control* | 36 | 0-17 | 5.2 | 3.5 | 83.3 |
| | | ds*MISO* | 7 | 1-11 | 4.6 | 3.0 | 100.0 |
